# Real-time three-dimensional MRI for the assessment of dynamic carpal instability

**DOI:** 10.1101/351593

**Authors:** Calvin B. Shaw, Brent H. Foster, Marissa Borgese, Robert D. Boutin, Cyrus Bateni, Christopher O. Bayne, Robert M. Szabo, Krishna S. Nayak, Abhijit J. Chaudhari

## Abstract

Carpal instability is defined as a condition where wrist motion or loading creates mechanical dysfunction, resulting in weakness, pain and decreased function. Often the diagnosis is made late when malalignment is visualized on static radiography, CT, or MRI. When conventional imaging methods do not identify the instability patterns, yet clinical signs associated with instability exist, the diagnosis of *dynamic instability* is often suggested to describe carpal derangement manifested only during the wrist’s active motion or stress. We addressed the question: can advanced MRI techniques provide quantitative measures for evaluating dynamic carpal instability and supplement standard static MRI acquisition? Our objectives were to [1] develop a real-time, three-dimensional MRI method to image the carpal joints during their active, uninterrupted motion; and [2] demonstrate feasibility of the method for assessing metrics relevant to dynamic carpal instability, thus overcoming limitations of conventional MRI. Twenty wrists (bilateral wrists of ten healthy participants) were scanned during radial-ulnar deviation and clenched-fist maneuvers. Images resulting from two real-time MRI pulse sequences, four sparse data acquisition schemes, and three constrained image reconstruction priors were compared. Image quality was assessed via blinded scoring by two radiologists and quantitative imaging metrics. Data acquisition employing sparse radial sampling with a gradient-recalled-echo acquisition and constrained iterative reconstruction (temporal resolution up to 135 ms per slice) appeared to provide a reasonable tradeoff between imaging speed and quality. This real-time MRI method effectively reduced streaking artifacts arising from data undersampling and enabled the derivation of quantitative measures pertinent to evaluating dynamic carpal instability.

## 1. Introduction

The wrist is comprised of a complex arrangement of carpal bones, ligaments, and tendons. These tissues collectively maintain biomechanical stability during physiologic motion and wrist loading^1–5^. When wrist ligaments are injured (e.g., owing to a fall or other excessive load on the wrist), the normal anatomical alignment of the carpal bones can be disrupted^6–8^, with consequent pain, dysfunction, and premature osteoarthritis^9^. The resulting joint instability will typically progress along a spectrum from early-stage “dynamic instability” (revealed only when the wrist is in motion or stress^10^; ^11^) to late-stage “static instability” (which can also be visualized with imaging even at rest)^12^. Thus, to assess dynamic instability, methods capable of capturing carpal bone kinematics during active motion or dynamic loading are needed. When physicians can diagnose dynamic instability early, interventions can be implemented to restore normal function^5; 10; 13; 14^

Magnetic Resonance Imaging (MRI) is the most commonly used tomographic imaging modality for evaluating ligament derangements in the wrist^15^. However, conventionally, MRI images the stationary, immobilized wrist and is therefore unable to directly evaluate dynamic carpal instability. Only a few studies have reported the use of fast MRI acquisition to visualize carpal kinematics^11; 16; 17^. This is primarily due to the slow encoding of k-space inherent to MRI that results in low temporal resolution (i.e., number of frames per second [fps])^18^, and the presence of susceptibility and other artifacts that may compromise image quality^11^. Several advances have been made recently to improve the performance of MRI for real-time data acquisition, such as using non-Cartesian sampling and/or sparse sampling coupled with constrained image reconstruction^18–22^. To date, these advances have been applied to investigate motion of the knee ^16; 23; 24^ and temporomandibular joint ^25; 26^ but not to carpal motion. For understanding, diagnosing and selecting appropriate treatment for dynamic instability of the carpus, there is increasing interest for dynamic imaging by hand surgeons and other clinicians ^27–29^. There has also been a concern that fast gradient-echo based pulse sequences that worked well for other joints may not be technically feasible in the moving wrist because magnetic field inhomogeneities generated by the significant displacement of the tissues during motion may generate substantial artifacts^11^. Therefore, we sought to develop a method for imaging the moving wrist, that overcame this challenge.

Our overall goal was to optimize real-time three-dimensional (3D) MRI (data acquisition and reconstruction) for imaging the unassisted, actively moving wrist and demonstrate the feasibility of deriving standardized imaging metrics relevant to assessing dynamic carpal instability. The total acquisition time was aimed to be quick (< 5 min) to have the ability to be incorporated into the current workflow of musculoskeletal radiology practice, as a supplement to a routine clinical MRI wrist scan. We compared two real-time MRI pulse sequences, four sparse data acquisition schemes, and three constrained image reconstruction priors, and then assessed these methods during the performance of wrist radial-ulnar deviation and the clenched-fist maneuvers.

## 2. Materials and Methods

### 2.A. Static Scanning and Comparison of Real-Time MRI Pulse Sequences

The study was performed on a 3.0T MRI system (Skyra, Siemens Healthcare, Erlangen, Germany) using a 32-channel radiofrequency (RF) head-coil (to accommodate the large range-of-motion of the wrist during its different maneuvers). The developed protocol consisted of two sections.

First, the gradient-recalled-echo (GRE)-based 3D volumetric interpolated breath-hold examination (VIBE) pulse sequence was optimized for high-spatial-resolution imaging (voxel size: 0.20 × 0.20 × 0.30 mm) of the immobilized wrist in the wrist’s neutral position. This scan served as our anatomical reference.

Next, two real-time MRI pulse sequences were implemented and optimized, namely (1) balanced steady-state free precession (bSSFP)-based, referred to by the vendor as true fast imaging with steady state precession (TrueFISP), and (2) fast GRE-based, referred to by the vendor as fast low angle shot (FLASH). The optimization was driven by determining a trade-off between the voxel size, bandwidth, slice thickness, field-of-view (FOV), and signal-to-noise-ratio (SNR), as described in subsequent paragraphs. Tissue contrast was chosen by tuning repetition time (TR), echo time (TE), and flip angle. Automated high-order B_0_ field shimming was performed while the wrist was held motionless in the neutral position before beginning the optimization procedures. Scan parameters were chosen by consensus between the authors.

Preliminary experiments with phantoms and two human participants were carried out to identify suitable scan parameters. The human participants lay in the “superman position” with one arm out-stretched above the head into the RF coil (standard position employed for clinical wrist MRI acquisition). The wrist was placed into an arm immobilizer and each participant was trained to perform the following two wrist maneuvers utilizing their full, active range-of-motion (absent pain) at a comfortable speed (i.e., continuously between the “start” and “stop” instruction interval of 20 s, completing at least 2 cycles of each maneuver): [i] radial-ulnar deviation, and [ii] the clenched fist maneuver with the wrist in the neutral position.

The scan parameters for bSSFP were matched with those reported in Boutin et al^11^. A comprehensive comparison of bSSFP and fast GRE was made by including Cartesian (rectilinear) and radial sampling schemes for each sequence. The key imaging parameters we converged on for both pulse sequences are given in Table 1. The other scan parameters common to both pulse sequences were field-of-view (FOV) of 120 mm^2^, acquisition matrix size of 112 x 112, in-plane resolution of 1.07 mm^2^, and slice thickness of 6 mm, with the 6 slices acquired in the coronal plane using the default sum-of-squares-based image reconstruction.

**Table-1:** Imaging parameters used for fast-GRE and bSSFP pulse sequences for scanning the actively moving wrist (R-radial, C-Cartesian).

| **Sequence** | **R-Fast-GRE** | **C-Fast GRE** | **R-bSSPF** | **C-bSSFP** |
| --- | --- | --- | --- | --- |
| Temporal Resolution (ms) | 315/194/134 | 351 | 476 | 477 |
| Voxel Size (mm<sup>3</sup>) | 1.07×1.07×6 | 1.07×1.07×6 | 1.07×1.07×6 | 1.07×1.07×6 |
| Radial Spokes or Cartesian Lines | 100/60/40 | 112 | 100 | 112 |
| TR/TE (ms) | [3.15/1.94/1.34]/1.74 | 3.13/1.74 | 4.77/1.76 | 4.25/1.75 |
| Bandwidth (Hz/pixel) | 990 | 990 | 930 | 930 |
| Flip Angle | 12 | 12 | 46 | 46 |

The total scan time for the static scan was 6 min, 30 s, while scans for each dynamic maneuver lasted 19 s (100 spokes), 12 s (60 spokes), and 8 s (40 spokes), keeping the same number of acquired time points (total of 10). Each maneuver was repeated thrice for test-retest analysis. At the end of each wrist scan, the raw k-space data were obtained from the scanner console. The optimized real-time sequences were compared for direct visualization of the actively moving wrist joints and for performance with respect to artifacts and image quality, as described below.

### 2.B. Temporal Resolution in Radial-fast GRE

We first confirmed that susceptibility/banding artifacts, that are unique to bSSFP sequences^11^ partially obscured the wrist during motion. Although dielectric pads could provide an option to reduce banding, the pads limit range of motion^11^ and therefore their use was avoided. The radially-sampled fast-GRE method provided reasonable spatial resolution with lesser artifacts (spokes=100) compared to bSSFP (discussed further in our Results); thus, we optimized the former further and analyzed the effect of voxel sizes and radial spokes on temporal resolution. Different in-plane spatial resolutions (by fixing the slice thickness to 6 mm), radial spokes (or angular undersampling), SNR, and their corresponding temporal resolutions were evaluated to determine suitable values relevant to this application. To satisfy the Nyquist criterion, 176 radial spokes are necessary to completely cover the k-space, 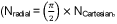 where N=number of frequency encoding lines). However, the resulting temporal resolution for 176 radial spokes was 554 ms (or less than 2 fps). We therefore chose 100 spokes as a starting point to have a moderate temporal resolution of about 3 fps, and then progressively reduced to acquiring 60 spokes (5 fps) and 40 spokes (7 fps) spokes.

### 2.C. Study Subjects

This study had approval from our Institutional Review Board (IRB) and written informed consent was obtained for each participant prior to study initiation. Three-dimensional (3D realtime MRI datasets were acquired from both wrists of ten healthy participants (6 men and 4 women, 27.6±6.1 years). Inclusion criteria included asymptomatic wrists, age <60 years, the ability to lie prone in the “superman position” during scanning, and the ability to follow directions to perform wrist motions while in the MRI scanner. Exclusion criteria were standard contraindications to MRI (including claustrophobia) and history of wrist pain and derangements (including trauma and arthritis). The static scan was conducted with the wrist in its neutral position, followed by dynamic scans, where the participants were asked to perform radial-ulnar deviation and clenched fist maneuvers. Each dynamic scan was repeated 3 times.

### 2.D. Constrained Image Reconstruction

Our approach to constrained image reconstruction using sparsity constraints to reduce undersampling artifacts was based on Lustig et al^18^. Accordingly, our image reconstruction problem was posed as:

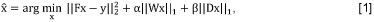

where is the reconstructed image, *F* and *y* are the Fourier encoding operator (undersampled) and the k-space measurements respectively, *W* and *D* denote the discrete cosine transform (DCT) and finite-differences (FD) operator respectively, and *α* and *β* are the regularization parameters that control the weighting for the two penalty functions. Sparsity is enforced by the choice of the norm for each of the *ℓ_1_*penalty functions. The DCT and FD schemes were applied only along the spatial dimension and not along the temporal dimension. In this framework, setting *α* = 0 or *β* = 0 results in only FD-sparsity or only DCT-sparsity-based minimization, respectively. The nonlinear conjugate gradient (CG) algorithm with backtracking line search was used to optimize the cost function in Eq. [1]. The gradient of the objective function was:

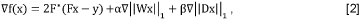

The ℓ_1_-norm is the sum of absolute values, so the function is not smooth. Therefore, we approximated the ℓ_1_-norm as 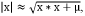 with a smoothing parameter, 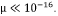 With this approximation, the derivative of the ℓ_1_-norm became 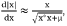 which was used to compute the gradient of the cost function in Eq. [1].

For the constrained reconstruction analysis, we compared the conventional gridding-based reconstruction, FD-sparsity (with α= 0), DCT-sparsity (with β = 0), and FD+DCT sparsity-based penalties. The non-linear CG algorithm was implemented using MATLAB (MathWorks, Natick, MA). Each channel’s data were individually reconstructed, and then the final image was combined by the root-sum-of-squares method. All four reconstruction methods were performed on a workstation with a 3.4 GHz Intel i7 processor and 128 GB RAM. Image Reconstruction Toolbox (IRT) ^30^ was used to perform the gridding-based reconstruction from radially sampled k-space data and to build the undersampled Fourier operator (F) to solve Eq. [1].

### 2.E. Parameter Tuning and Convergence

Choosing an appropriate regularization parameter (α or β in Eq. [1]) is crucial for suppression of undersampling artifacts in reconstruction. To compare regularization schemes, we adopted the L-curve approach, which eliminates the heuristic selection of the regularization parameter(s). This requires an iterative search by solving the minimization schemes with various regularization parameters and plotting the model error 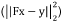 and prior error 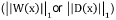 to pick the elbow of the curve that denotes the optimal regularization parameter. Note that selection of α and β simultaneously for FD+DCT sparsity optimization is not straightforward using L-curve as it requires determining the corner of the hypersurface^31^. Hence, both α and β were empirically chosen for this estimation. Further, we investigated the convergence behavior of the three types of penalties to evaluate the consequence of using more than one penalty in the cost function. The gridding reconstruction was set as the initial guess for all the three iterative reconstructions.

### 2.F. Image Quality Assessment

Two fellowship-trained, board-certified musculoskeletal radiologists, one with 21 years and the other with 7 years of experience, participated in the human observer study. They first met with the researchers together, finalized the viewing settings, and reached consensus on scoring images from the different reconstructions of the wrist radial-ulnar deviation maneuver based on a 5-point scale (4: strongly agree; 3: agree; 2: neutral; 1: disagree; 0: strongly disagree) in response to three clinically-relevant statements:

1. *I can confidently measure the scapholunate (SL) gap.* This statement is of relevance for assessing dynamic SL instability^32^;
2. *I can confidently measure the scaphoid translation with respect to distal radius.* This statement is of value to understanding scaphoid-radial motion and maltracking after trauma^33^;
3. *I can confidently measure the trapezium translation with respect to the scaphoid.* This statement is of importance in determining dynamic instability associated with the scaphoid-trapezium joint^34^.

For the evaluation study, each radiologist independently scored a total of 120 images (10 right wrists x 4 reconstructions (gridding, FD-sparsity, DCT-sparsity, and DF+DCT sparsity) x 3 sampling schemes (100, 60, and 40 spokes)). They were blinded to the type of reconstruction and sampling scheme, and the real-time 3D datasets were presented in a random fashion. The overall performance of each reconstruction scheme was quantified as the scores from one question for a given reconstruction method, averaged (mean and standard deviation) across different number of spokes. The inter- and intra-rater reliability was estimated using Cohen’s kappa (κ) coefficient for the same four reconstruction methods. The intra-rater reliability was evaluated by asking the one of the radiologists to repeat the scoring of randomly chosen 36 images (four subjects). The interval between the two scoring sessions was about a month.

Measures of image quality were also obtained from our data in a test-retest setting. For this, Bland-Altman analysis was performed to quantify test-retest capability of the radial-fast-GRE sequence for varying number of spokes. The plots were computed based on SNR estimated from two measurements chosen randomly from the three repeated measurements. SNR was defined as the ratio of mean signal intensity in the region of interest (ROI) containing the carpal bones to the standard deviation of the background signal. The x- and y-axes of the Bland-Altman analysis were defined as the average and difference, respectively, in SNR at a single time point for the first and second repeat respectively during radial-ulnar deviation maneuver for each participant.

### 2.G. Scapholunate (SL) Gap Measurement

To demonstrate the application of the proposed real-time MRI acquisition for providing quantitative measures of commonly presented SL injuries, we measured the SL gap of all ten subjects’ right and left wrists during the radial-ulnar deviation and clenched-fist maneuvers. Single slices obtained from gridding reconstruction (spokes=100, i.e., lowest temporal resolution, and spokes=40, i.e. highest temporal resolution) and the FD+DCT-sparsity reconstruction (spokes=40, i.e. highest temporal resolution) were used to assess this gap, measured as the distance between the midpoints of the scaphoid and lunate articulating surfaces at the SL joint, using the ‘ruler’ tool in ImageJ (National Institutes of Health, Bethesda, MD).

## 3 Results

### 3.A. Comparison of bSSFP and fast-GRE acquisitions

Figure 1 shows the coronal slices for comparing the real-time MRI pulse sequences during the wrist radial-ulnar deviation maneuver. In each case, the number of spokes or Cartesian lines in k-space were the same (100 lines) and images were reconstructed by the gridding-method. Figure 1 also indicates the wrap-around artifact, which is an inherent artifact associated with Cartesian sampling, and the characteristic banding artifacts arising from the bSSFP pulse sequence^11^. Radial sampling was found to be more robust compared to Cartesian sampling when dealing with these artifacts. Based on these assessments, the radially-sampled-fast-GRE approach was optimized further.

**Fig 1:**
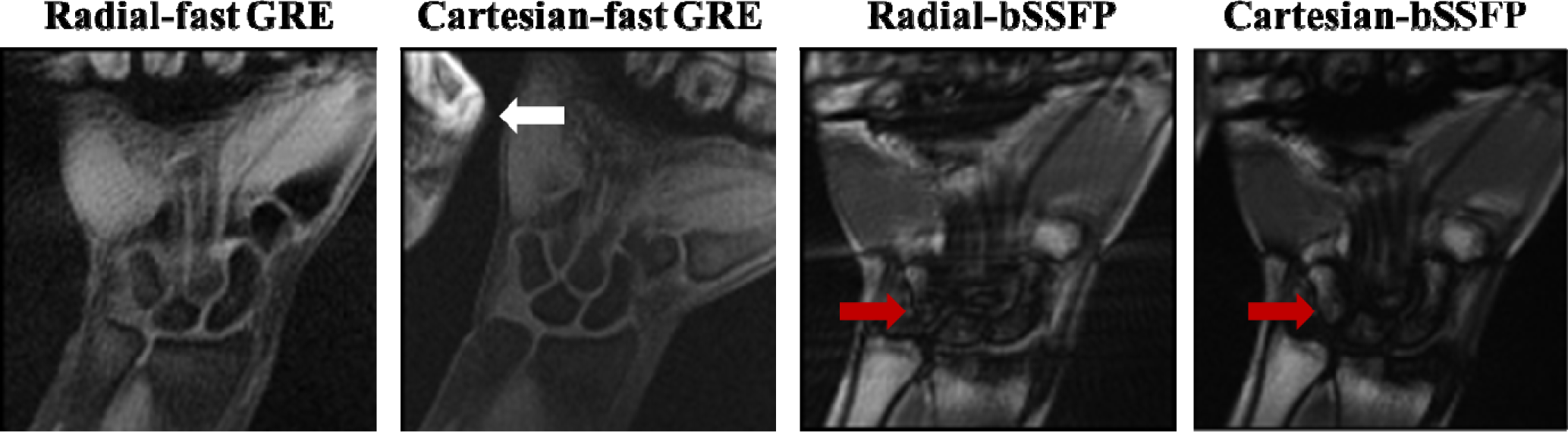
Comparison of real-time MRI pulse sequences, namely radial-fast GRE, Cartesian-fast GRE, radial-bSSFP, and Cartesian-bSSFP from healthy, asymptomatic study participants during radial-ulnar deviation. White and red arrows point to the wraparound effect and banding artifacts, respectively.

### 3.B. Temporal Resolution in Radial-Fast-GRE

Figure 2 illustrates the trade-off between temporal resolution, number of spokes, and pixel size (in-plane resolution) for the radial-fast-GRE pulse sequence. Reducing the number of spokes (angular undersampling) resulted in a higher temporal resolution compared to just increasing the pixel size. Our assessments converged on a pixel size of 1.1 mm^2^ and the parameters in Table 1, as a trade-off between SNR and joint space delineation.

**Fig 2:**
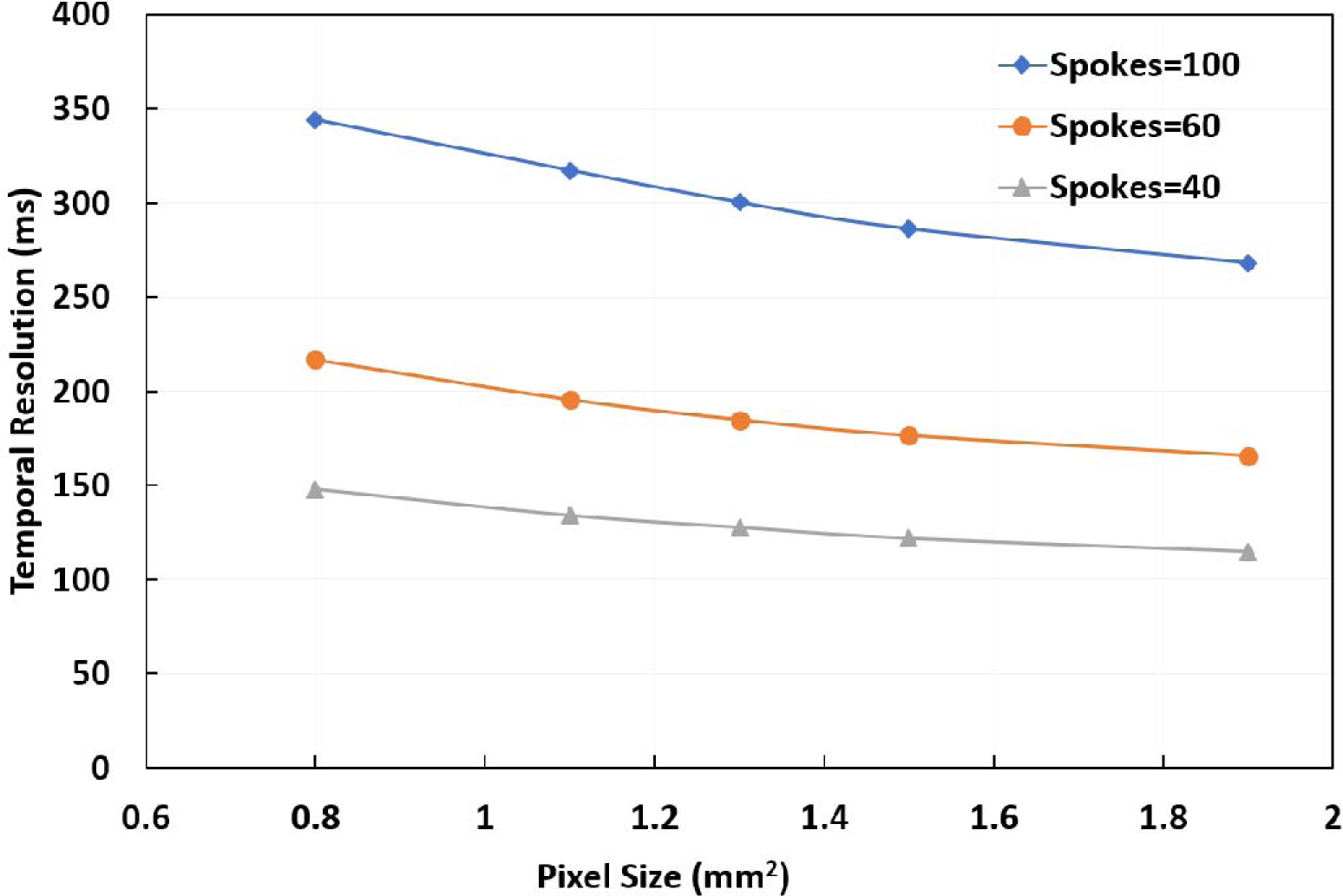
Illustration of different temporal and pixel sizes achievable with the radial-fast GRE sequence given in Table 1. The graph also describes the trade-off between number of spokes and temporal resolution.

### 3.C. Constrained Image Reconstruction

The best regularization parameters estimated through L-curves were used for DCT-sparsity and FD-sparsity-based reconstruction, as the values at the maximum curvature (independently for 100, 60, and 40 spokes). For the FD+DCT-based sparsity penalty, the regularization parameters were empirically determined such that the range of *α* was found to be [1×10^−5^: 1×10^−6^] and *β* was found to be [1×10^−6^: 1×10^−8^]. These values provided a fair balance between priors (for spokes=100, 60, and 40). In addition, we found that with higher undersampling, higher values of α and β were required to reduce the undersampling artifacts. Figure 3 compares the convergence curves of DCT-sparsity, FD-sparsity and FD+DCT-sparsity-based penalties (corresponding to spokes=40 from Figure 4), our fastest method for data acquisition. The curves show the number of iterations required by each type of penalty to reach an empirically chosen stopping criterion of 1×10^−6^ (i.e., when the relative error of the cost function was less than 1×10^−6^). Note that the FD+DCT-sparsity penalty converged faster than the other two penalties indicating the improvement via using multiple priors in the cost function (Eq. [1]).

**Fig 3:**
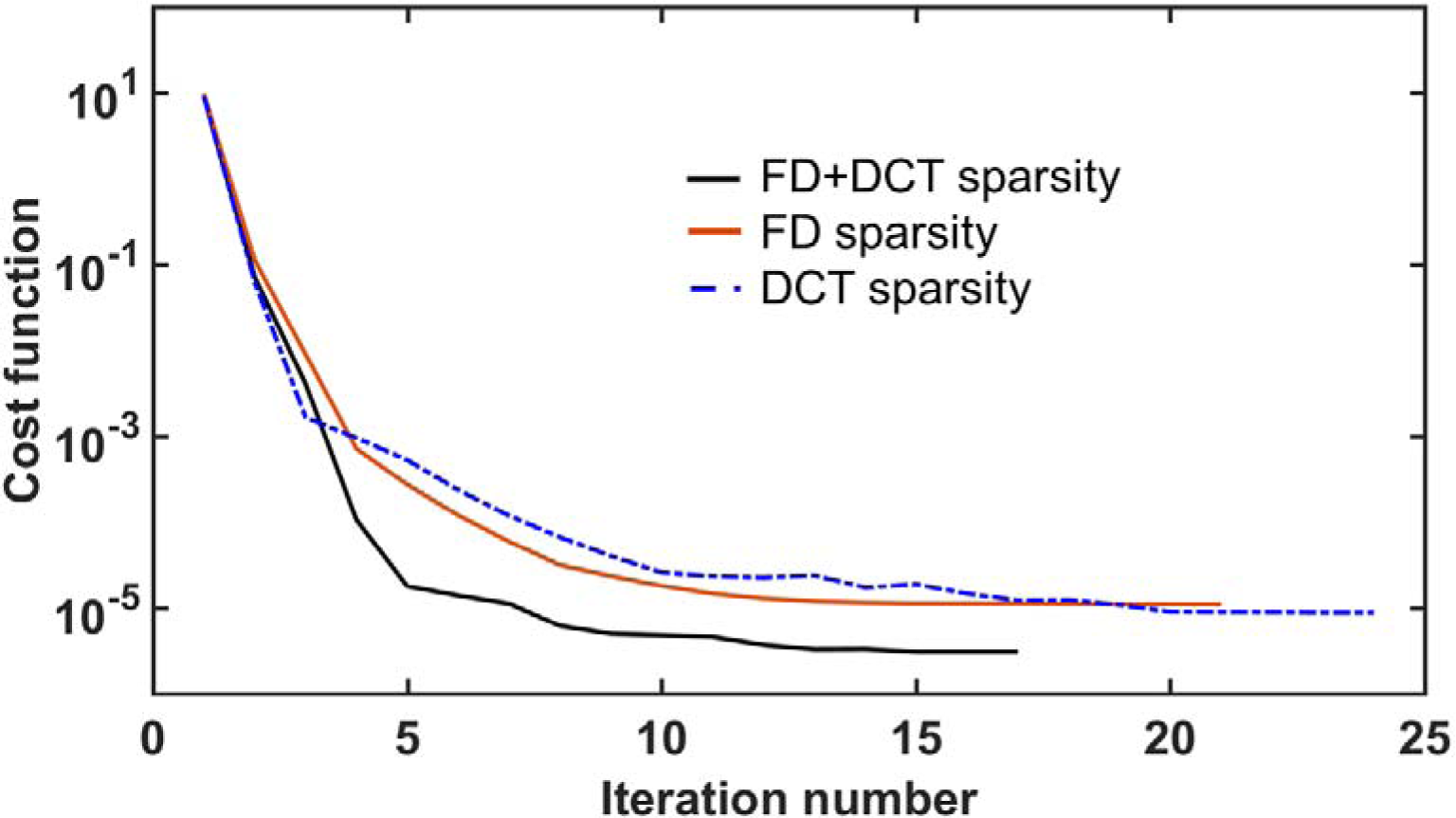
Graph illustrating the convergence curves for FD+DCT sparsity, FD-sparsity, and DCT-sparsity-based penalties for the case of 40 spokes in Fig. 4.

The reconstructed coronal slices corresponding to a single time point from the reconstruction schemes during radial-ulnar deviation and clenched-fist maneuvers of the wrist are shown in Figures 4 and 5 respectively. Comparison of different reconstruction schemes revealed that all iterative reconstruction schemes were effective in reducing the overall image noise apparent in gridding reconstructions. Although the FD-sparsity-based penalty reduced streaking artifacts, it also induced spatial blurring. This can be attributed to the staircasing artifact associated with the first-order penalty and that the regularization parameter estimated by the L-curve method may not be optimal from a visualization point of view. The DCT-sparsity-based penalty preserved anatomical features, such as edges, but still had some residual streaks in the reconstruction (evident with spokes=40). These artifacts were mitigated by simultaneous incorporation of FD and DCT-sparsity-based penalties, especially with lower number of spokes (60 and 40), by rendering much sharper anatomical information. Representative movies showing radial-ulnar deviation (corresponding to Figure 4) and clenched fist maneuvers (corresponding to Figure 5) are available as Supplemental Movies M1 and M2, while snapshots over time are shown in Supplemental Figure 1. Images reconstructed from 100 spokes rendered a discontinuous motion while images corresponding to 60 and 40 spokes rendered a smoother motion. The sharpness for feature edges obtained with FD+DCT-sparsity-based reconstruction showed that constrained reconstruction can improve the visualization of the joint spaces in the wrist particularly with 60 and 40 spokes. The FD+DCT-sparsity-based penalty offered an overall better visualization of the joint spaces compared to gridding, FD-sparsity or DCT-sparsity-based methods.

**Fig 4:**
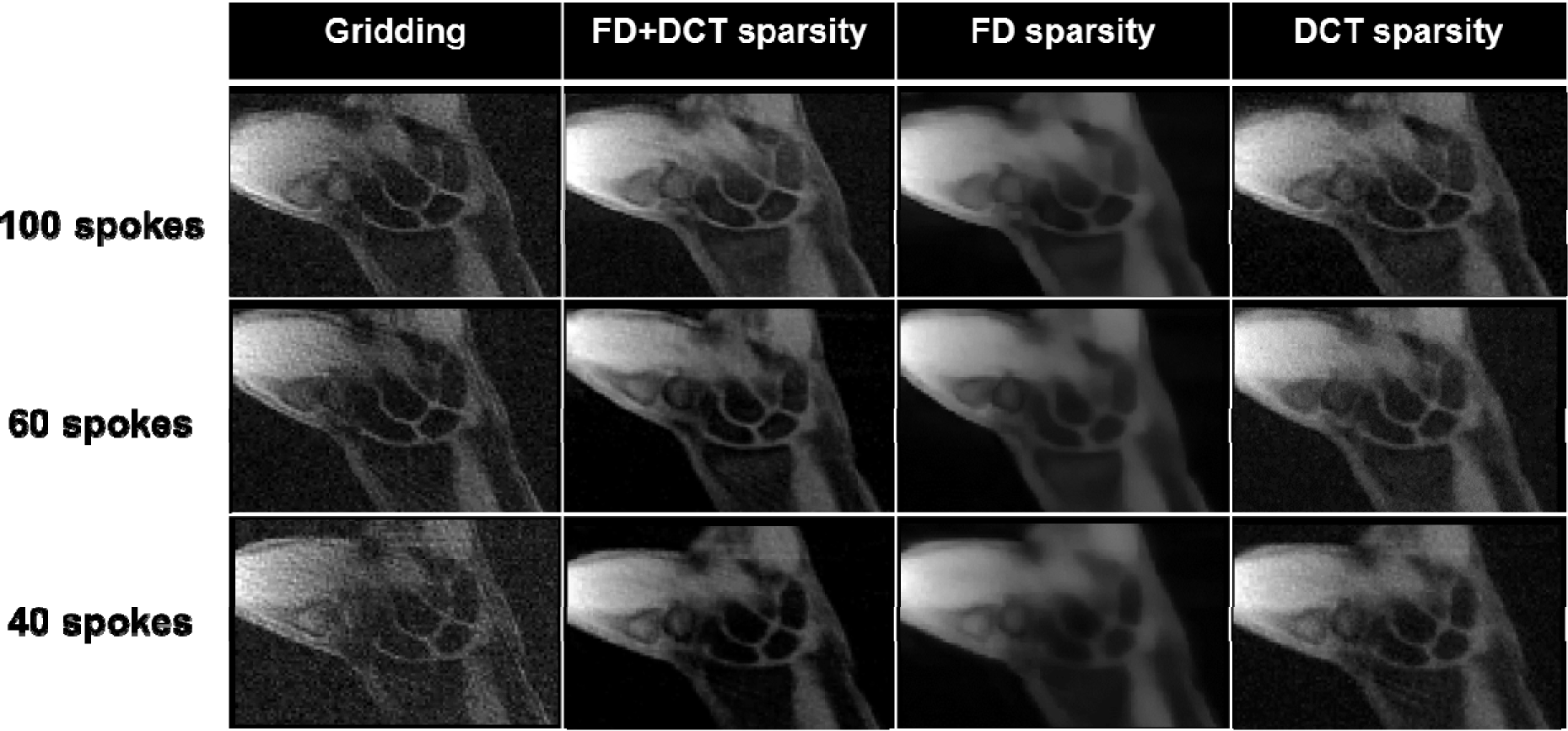
Representative coronal slices of a participant’s right wrist at a time point during radialulnar deviation obtained from gridding, FD+DCT sparsity, FD-sparsity, and DCT-sparsity-based reconstruction schemes using 100, 60, and 40 spokes.

**Fig 5:**
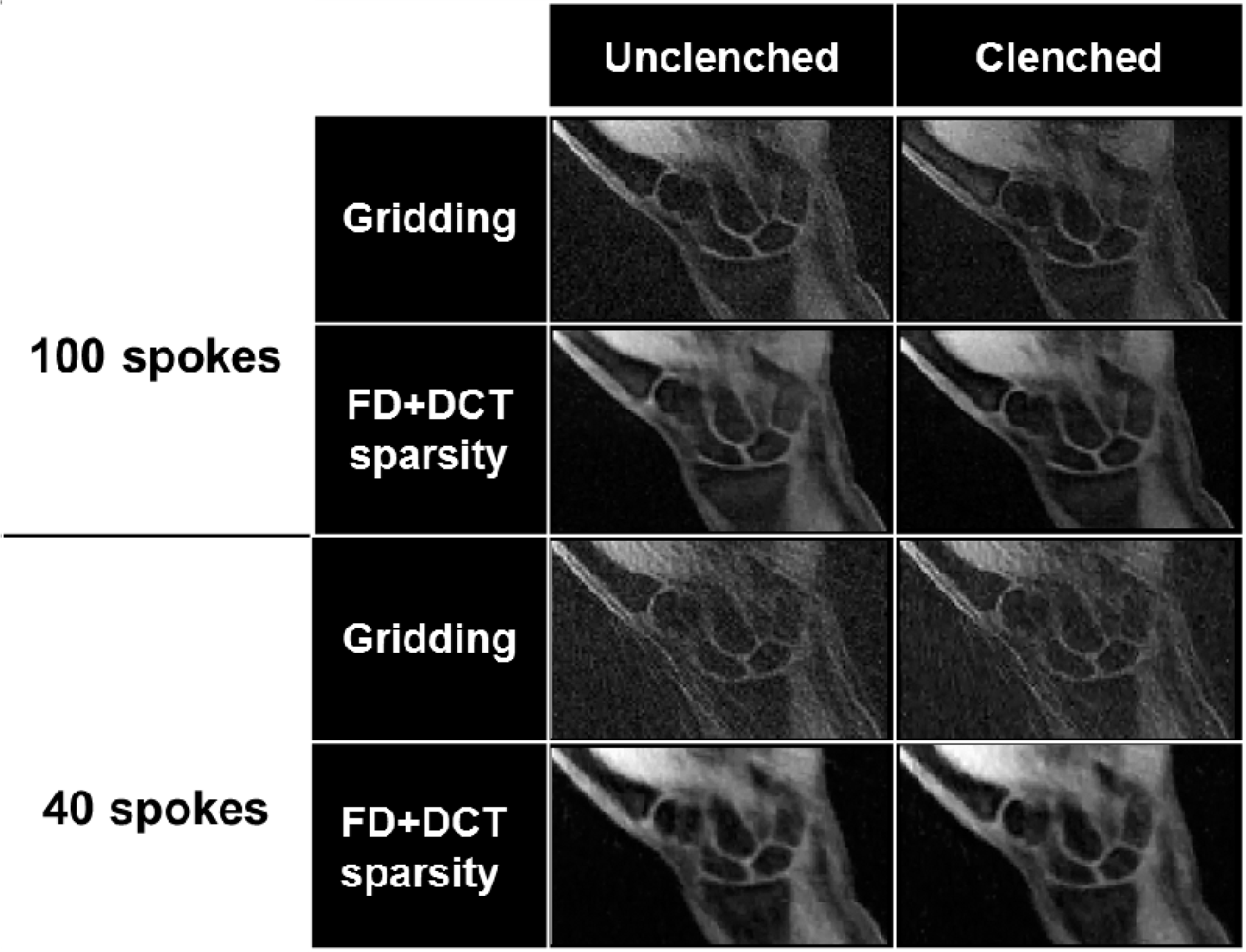
The same participant’s right wrist as shown in Fig. 4 at a time point during clench-unclench maneuver using the gridding, FD+DCT sparsity, FD-sparsity, and DCT-sparsity-based reconstruction schemes using 100, 60, and 40 spokes.

### 3.D. Image Scoring by Radiologists

The averaged (mean ± standard deviation) visual scores assigned by the two radiologists for all the reconstruction schemes are given in Table 2. The average scores suggest that the FD+DCT-sparsity-based reconstruction scheme consistently performed better than other schemes for all three clinical questions. The intra-rater *κ* had a range between 0.5 and 0.7 for the various reconstruction schemes and spokes (highest for FD+DCT-sparsity), indicating moderate to substantial intra-rater reliability.

**Table-2:** Table-2: Average image scores of ten subjects’ datasets for all four reconstruction schemes by the two radiologists (R1 and R2) with clinical questions. Values given are mean±standard deviation (SD) of radiologist-assigned image quality scores. The statements were: S1: I can confidently measure the SL gap; S2: I can confidently measure scaphoid translation with respect to distal radius; S3: I can confidently measure trapezium translation with respect to the scaphoid.

| **Clinical**<br>**Score** | **Gridding** |  | **FD+DCT sparsity** |  | **DCT sparsity** |  | **FD sparsity** |  |
| --- | --- | --- | --- | --- | --- | --- | --- | --- |
|  | R1 | R2 | R1 | R2 | R1 | R2 | R1 | R2 |
| S1 | 1.3±1.0 | 0.6±0.6 | 3.0±0.9 | 1.7±1.0 | 2.8±1.0 | 1.4±0.7 | 2.7±0.9 | 1.2±0.8 |
| S2 | 1.6±1.0 | 0.6±0.7 | 3.2±0.9 | 1.9±0.9 | 3.0±1.0 | 1.8±0.9 | 3.2±0.9 | 1.4±1.0 |
| S3 | 2.0±1.3 | 0.7±0.7 | 3.1±1.0 | 1.8±1.0 | 3.1±0.9 | 1.6±1.0 | 3.1±1.0 | 1.0±0.8 |

### 3.E. SL Gap Measurement

The SL gap was measured for gridding-based reconstruction (100 and 40 spokes) and the FD+DCT-sparsity-based method (40 spokes). Figure 6 shows the mean and standard deviation of SL gap across time during radial-ulnar deviation, while Table 3 shows quantitative values for clenched-fist and unclench maneuvers. On comparing the measurements obtained from gridding (100 and 40 spokes) and FD+DCT-sparsity (40 spokes), we observed that the performance of the FD+DCT-sparsity-based method was closer to gridding with 100 spokes than gridding with 40 spokes. This was predominantly attributed to streaking artifacts (in gridding with 40 spokes). These values are within the range of values reported in the literature^11; 35^. The SL gap widened, albeit insignificantly, as the wrist went from the relaxed to the clenched-fist position, also consistent with the literature^11; 36^.

**Fig 6:**
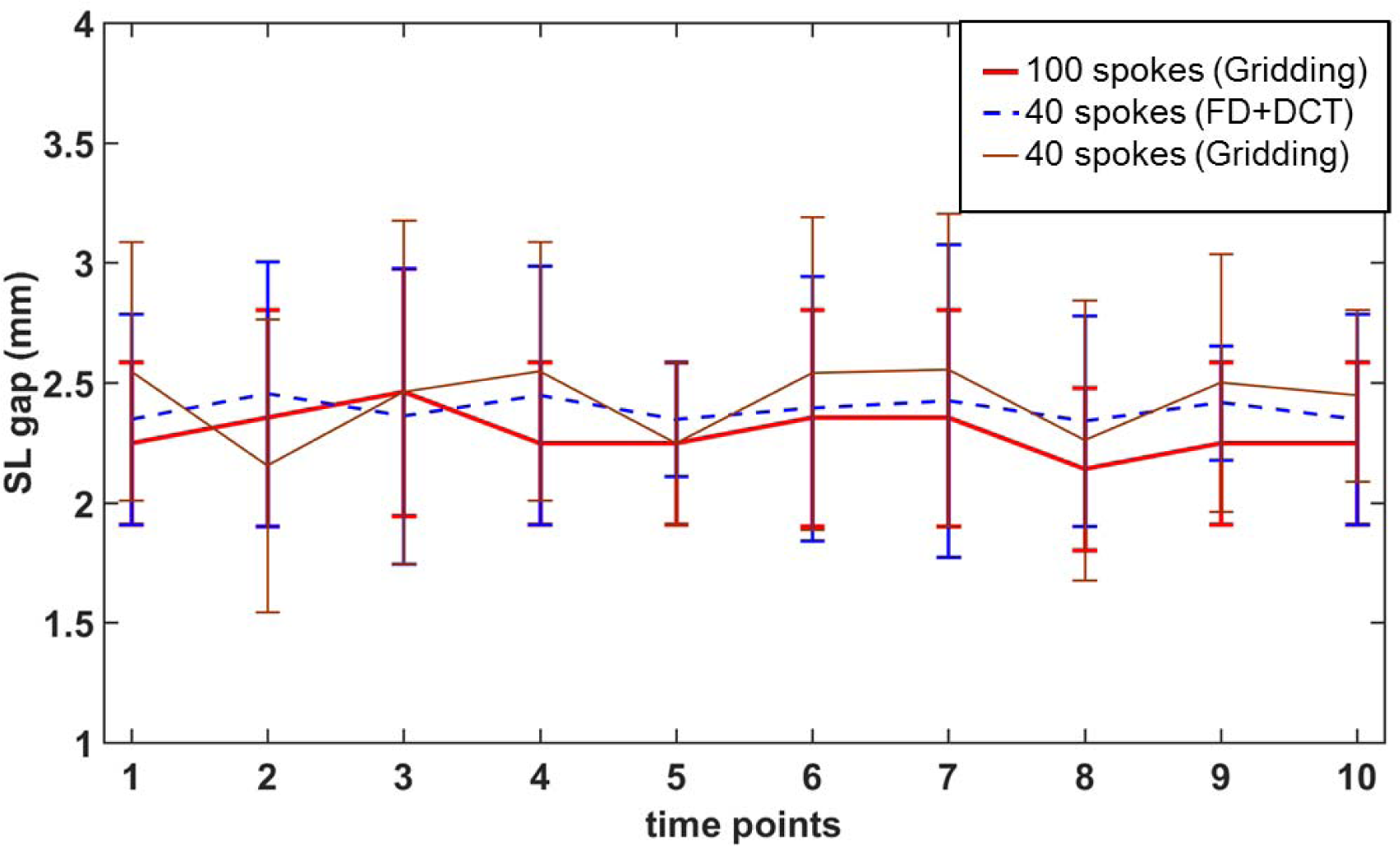
The SL gap (mean ± standard deviation (SD)) during radial-ulnar deviation maneuver at every time point measured by tracking a single slice through the wrist for the 10 subjects. Values from only the right wrist are shown for clarity. The measurements were obtained from gridding-based reconstruction for 100 and 40 spokes, and DCT+FD sparsity-based reconstruction for 40 spokes.

**Table-3:** Table-3: SL gap measurement during the clenched-fist maneuver for all ten subjects. Values are given as mean±SD.

| Metric | Unclenched fist |  |  | Clenched fist |  |  |
| --- | --- | --- | --- | --- | --- | --- |
|  | Gridding<br>Spokes=100 | FD+DCT<br>Spokes=40 | Gridding<br>Spokes=40 | Gridding<br>Spokes=100 | FD+DCT<br>Spokes=40 | Gridding<br>Spokes=40 |
| SL gap (mm) | 2.3±0.3 | 2.4±0.5 | 2.5±0.5 | 2.4±0.5 | 2.5±0.7 | 2.3±0.6 |

### 3.F. Test-Retest Analysis

Bland-Altman analysis used to quantify test-retest capability of radial-fast GRE sequence for spokes=100, 60, and 40 is shown in Figure 7. The plot shows that most of the SNR points lie within the 95% confidence interval of the mean of the differences, quantifying the robustness of radial-fast-GRE sequence. Also note the decreasing SNR values on the x-axis with spokes=100, 60, and 40, suggesting that reducing the spokes from 100 to 40 reduces SNR by almost half, as would be expected.

**Fig 7:**
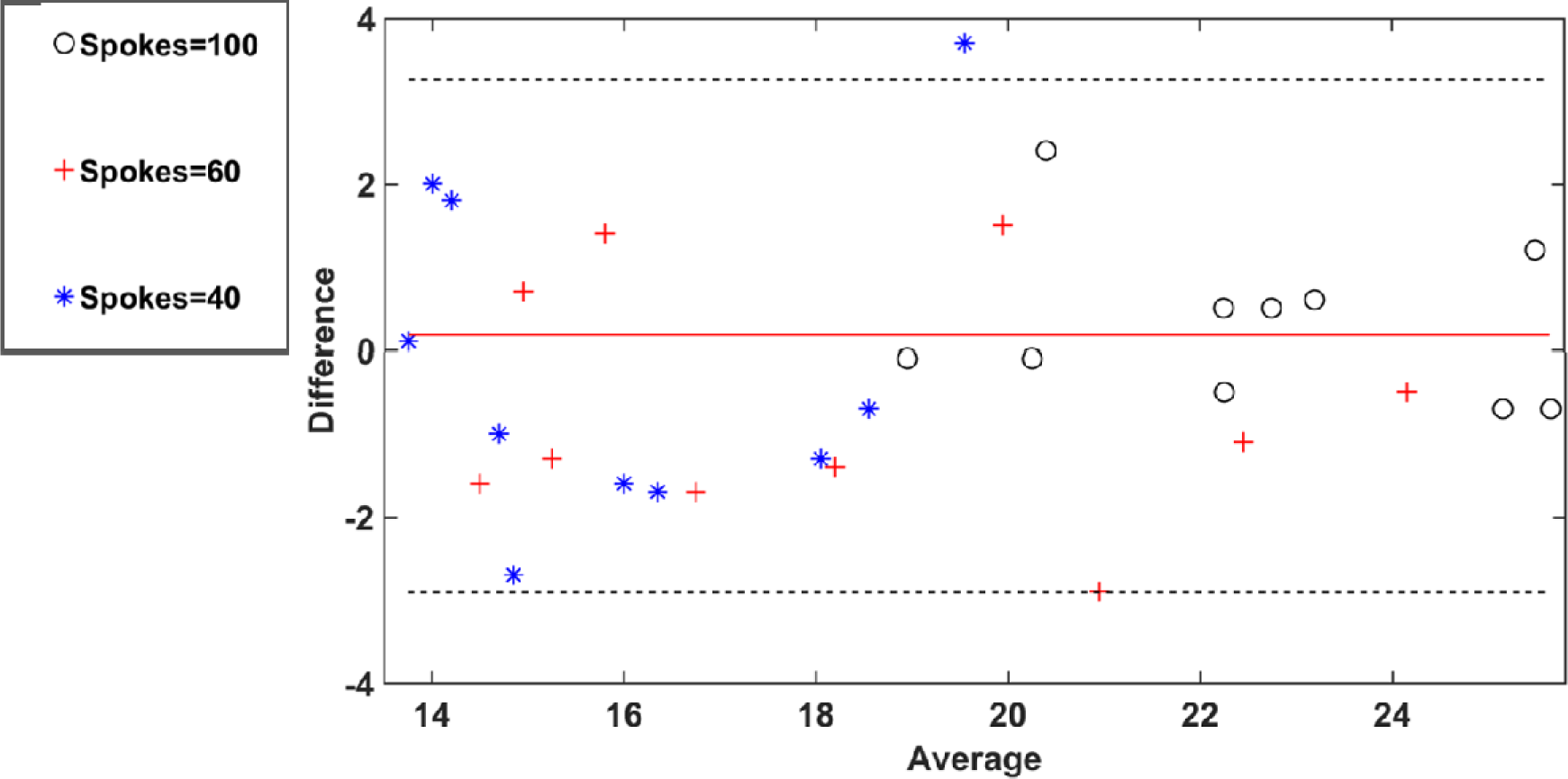
The Bland-Altman plot to quantify test-retest capability of radial-fast GRE sequence for spokes=100, 60, and 40. The y- and x-axes indicate the difference in SNR between first and second repeat and average of the SNR’s between first and second repeat respectively during radial-ulnar deviation for all the ten subjects’ at a time point. Values from only the right wrist are shown for clarity. The red line indicates the mean of the differences, while the 95% confidence intervals are denoted by the pair of dotted black lines.

## Discussion

Clinical MRI protocols are currently limited to assessing the wrist in its static position. However, dynamic carpal instability is only manifested and diagnosed during normal wrist motion or with loading. In Boutin et. al.^11^, the achievable temporal resolution of dynamic wrist imaging was about 400-600 ms per slice using a 2D bSSFP sequence. Additionally, since bSSFP is susceptible to banding artifacts, a dielectric pad containing a perfluorocarbon liquid was attached to the wrist to reduce the magnetic field inhomogeneity, but it had the undesired effect of limiting the range of motion during some maneuvers^11^. The use of the fast-GRE sequence, which utilizes shorter TE and TR, is less sensitive to tissue susceptibility differences, making it more suitable for realtime imaging. We used a radial-sampling scheme, instead of Cartesian sampling, as the former is less sensitive to motion artifact/temporal blurring due to inherent averaging of low spatial frequencies ^37^. This work shows an approximately 4-fold increase in temporal resolution compared to existing methods, i.e. 4 additional slices (either in space or time) could be acquired via our proposed method in the time required for acquiring data for a single slice with previous methods^11, 16^.

The radiologist-provided image quality scores suggest that FD+DCT-sparsity-based penalty significantly improved the reconstructed image quality (by about two-fold) in comparison with gridding method. Use of multiple sparsity-based priors, such as FD+DCT-sparsity, was shown to have fewer overall artifacts due to undersampled data compared to single priors such as FD or DCT, consistent with other applications^38, 39^. Since the wrist images are not inherently sparse in their pixel representation, we performed preliminary evaluation of two sparsifying transformations, DCT and the Daubechies-4 wavelet coefficients. We did not find significant differences between the two transformations in terms of the reconstruction performance. Therefore, we chose to use DCT as a sparsifying transformation in this work. Wavelets, such as Daubechies 4, have the restricted isometry property that makes them more suited for compressive sensing^18^. A detailed comparison of DCT versus wavelets will be conducted in future work.

Dynamic computed tomography (CT) has been successfully used to investigate the normal kinematics and pathokinematics of the actively moving wrist^40–42^ and to provide measures for assessing dynamic carpal instability^12^. However, CT involves ionization radiation, has poor soft tissue contrast compared to MRI, and generally is ordered as a separate exam. We aimed to design our real-time protocol as a supplement to a standard static MRI exam, which avoids the additional time, expenses, and risks associated with dynamic CT. A comparison of real-time MRI with dynamic CT was outside the scope of this work but will be pursued in the future.

The radiologists’ scores support the utility of the real-time datasets for assessing wrist features of interest in dynamic carpal instability. Of particular note, we used the real-time images to measure the SL gap, which can be used to identify pathologic changes in carpal alignment in patients with dynamic SL instability. These changes in the SL gap showed that real-time MRI may provide information beyond that from a static MRI examination for cases of suspected dynamic carpal instability. In addition, real-time MRI of the wrist could serve as a valuable imaging tool for determining improved treatment plans and monitoring response to therapy, e.g., stem cell or prolotherapy for SL tear^43^. To the best of our knowledge, this is the first work that has demonstrated the use of radial-fast-GRE sequence with constrained reconstruction to investigate the kinematics of the moving wrist.

Our study also had limitations. First, the constrained, iterative (offline) reconstruction scheme was computationally intensive compared to the gridding (online) reconstruction scheme. The average reconstruction time for FD+DCT-sparsity reconstruction was about 80 s per slice, while the gridding scheme required only 15 s. This limitation can be addressed in the future by using high-performance computing techniques like graphics processing units (GPUs) to accelerate image reconstruction^41^. Second, the regularization scheme was applied only in the spatial dimension and not along the temporal dimension. Application of temporal regularization schemes could further provide improved reconstruction quality by reducing temporal blurring. Simultaneous use of spatial regularization and temporal regularization (such as temporal FD, and nuclear norm) could provide favorable image quality (reduced temporal blurring) with fewer radial spokes (< 40), thereby improving the temporal resolution^44; 45^. Future work will focus on unifying reconstruction and motion estimation for k-space. In addition, non-Cartesian sampling schemes suitable for motion imaging, such as golden-angle and spiral sampling, and image reconstruction methods utilizing machine learning approaches will be explored in our future work. These strategies may be effective to reconstruct artifact-free wrist images with a higher spatial and temporal resolution, allowing more subtle SL gap differences to be measure with a higher accuracy.

## Conclusion

We proposed a real-time MRI scan protocol with a temporal resolution of 135 ms per slice (~7 fps) that can be used to assess continuous, active, and uninterrupted wrist motion, and extract anatomical information relevant to the diagnosis of dynamic carpal instability. The proposed real-time protocol utilizes a short scan time, typically less than 20 s per maneuver, which means it can be incorporated as a supplement to a standard clinical wrist MRI exam (typically 30-45 min long) with minimal disruptions to clinical workflow. The quick supplemental technique to examine the wrist kinematics could represent a paradigm shift for clinical MRI and further our understanding of baseline patterns of carpal kinematics, and improve our ability to assess dynamic carpal instability.

## Author Contributions Statement

All authors have reviewed, provided comments on, and approved of the paper prior to submission for publication. Conceived and designed the experiments: CBS, BHF, RDB, COB, RMS and AJC. Performed the experiments: CBS, BHF, RDB, COB, and AJC. Analyzed the data: CBS, BHF, RDB, COB, and AJC. Contributed to analysis tools and data interpretation: COB, RMS, KSN. Wrote and contributed to the paper: All authors.

## Acknowledgements

This study was funded by the National Institutes of Health (NIH) grants K12 HD051958 and R03 EB015099, and National Science Foundation (NSF) GRFP Grant No. 1650042. The views expressed in this article are the authors’ own and do not necessarily represent the views of the NIH or NSF. The authors would like to acknowledge the contributions of Drs. Michael H. Buonocore, John C. Hunter, Eva Escobedo, and Costin Tanase, and John Brock and Gerard Sonico, all from the University of California Davis.

## Supplemental Data

Supplemental Movie 1: Fly-through from the real-time MRI in time and space (slices) during the performance of the radial-ulnar deviation maneuver for FD+DCT-based constrained reconstruction with spokes=40.

Supplemental Movie 2: Fly-through from the real-time MRI in time and space (slices) during the performance of the clenched-fist maneuver for FD+DCT-based constrained reconstruction with spokes=40.

**Supplemental Figure 1:**
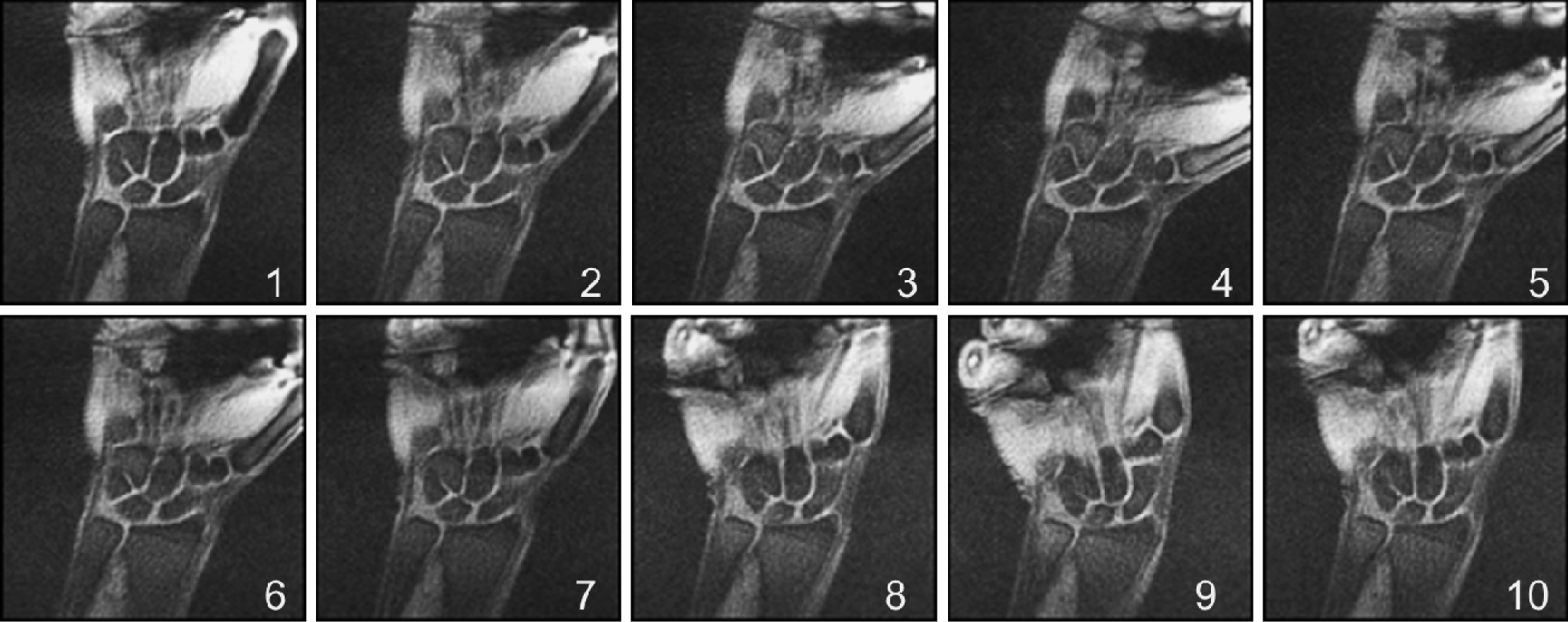
Representative coronal section through the wrist at the various time-points during the radial-ulnar deviation maneuver.

